# Methyl-CpG2-binding protein 2 mediates overlapping mechanisms across brain disorders

**DOI:** 10.1101/819573

**Authors:** Snow Bach, Niamh M. Ryan, Paolo Guasoni, Aiden Corvin, Daniela Tropea

## Abstract

Methyl-CpG binding protein 2 (MeCP2) is a chromatin-binding protein and a modulator of gene expression. Initially identified as an oncogene, *MECP2* is now mostly associated to Rett Syndrome, a neurodevelopmental condition, though there is evidence of its influence in other brain disorders.

We design a procedure that considers several binding properties of MeCP2 and we screen for potential targets across neurological and neuropsychiatric conditions.

We find MeCP2 target genes associated to a range of disorders, including - among others-Alzheimer Disease, Autism, Attention Deficit Hyperactivity Disorder and Multiple Sclerosis. The analysis of biological mechanisms and pathways modulated by MeCP2’s target genes shows that such mechanisms are involved in three main processes: neuronal transmission, immuno-reactivity and development.

These results suggest that similar symptoms present in different pathologies have a common molecular basis, and that treatments for one condition have potential applications to related disorders.

## Introduction

MeCP2 is a protein that binds methylated CpG dinucleotides to modify chromatin and control gene expression. Although MeCP2 acts mainly as repressor of transcription, it also plays a role as an activator through interactions with other co-factors (CREB1).

Besides direct DNA binding, MeCP2 controls gene expression through other mechanisms such as splicing and regulation of miRNA. MeCP2 interacts with splicing proteins and can recognise exon methylation to promote inclusion of alternatively spliced exons. It also affects gene expression indirectly by regulating miRNA or DGCR8 miRNA processing protein, with consequences for neurogenesis and differentiation (for an overview of MeCP2 protein and function, see a review by Guy et al., 2011).

Initially identified as an oncogene, *MECP2* mutations have been detected in several neurodevelopmental disorders of monogenic and polygenic aetiology. The *MECP2* gene is mostly associated with Rett Syndrome (RTT): a progressive X-linked neurological disorder that primarily affects females. However, mutations in *MECP2* are present in other conditions, and indeed similar phenotypes have been identified across disorders such as Angelman Syndrome, Tuberous Sclerosis, Fragile X and Autism Spectrum Disorders (ASD).

MeCP2 deficiency in RTT has been shown to indirectly downregulate UBE3A - the causative gene in Angelman Syndrome. The interaction occurs through lack of regulatory repression on the maternal imprinting centre (Makedonski et al., 2005), with the de-repression of the *UBE3A* antisense gene and inhibition of UBE3A synthesis. In Fragile X Syndrome, caused by silencing of *FMR1*, MeCP2 has been reported to be involved in epimutations of *FMR1* (Coffee et al., 2002). Patients with Tuberous Sclerosis display similar phenotypes to patients with RTT, both presenting with intellectual disability, epilepsy, autistic traits and abnormal neurological phenotypes. In Tuberous Sclerosis, the genes *TSC1* and *TSC2* are mutated with a subsequent alteration of hamartin and tuberin tumour suppressor proteins, respectively (Khwaja and Sahin, 2011). MeCP2 represses both TSC1 and TSC2 and promotes proliferation and differentiation in human embryo lung fibroblast cells (Wang et al., 2017). There is a strong association between RTT and ASD: in fact, RTT was originally classified as part of ASD. Autism is broadly characterised by abnormal social behaviour, stereotypies, language use and impaired joint attention to varying degrees which is also seen in some phases of RTT, thereby suggesting shared interplay. Most ASD displays complex genetic aetiology with multiple genes, and mutations in *MECP2* have been reported in patients with autism (Li et al., 2005). MeCP2 is also involved in several neuropsychiatric and neurological disorders and its dysregulation can have functional consequences (reviewed by Ausió et al., 2014).

Considering the direct and indirect scope of MeCP2 function, we hypothesize that MeCP2 is involved in the regulation of genes associated with a range of neurological and neuropsychiatric disorders. We develop a procedure to identify potential MeCP2 binding sites over false positives and we apply this procedure to selected gene-sets derived from several studies involving neuropsychiatric and neurological disorders. The gene sets analysed include GWAS, and meta-analyses for several brain disorders: schizophrenia (SCZ), ASD, attention deficit hyperactivity disorder (ADHD), depression, anorexia, epilepsy, Alzheimer disease (AD), Parkinson disease (PD), Huntington’s disease, amyotrophic lateral sclerosis (ALS) and multiple sclerosis (MS).

Using this approach, we show that MeCP2 binds the promoters of genes associated with brain disorders more often than expected by chance. We identify genes that are potential targets of MeCP2 and use these genes to perform Protein-Protein Interaction (PPI), Gene Ontology, and pathway enrichment analysis for each disorder. Our results reveal unexpected connections across different pathologies and suggest that common molecular mechanisms are active across several brain disorders.

## Results

### Establishing a High Affinity MeCP2 Binding Procedure

To identify candidate target genes of MeCP2, we establish a procedure that considers both nucleotide sequences and the content of GC in promoters. This procedure implements a PWM used to identify MeCP2 preferred sequences (Klose et al., 2005) (Fig. 1a), combined with GC content (in percentage) of the promoter sequence, and we establish the optimal range of the selected variables.

**Fig. 1.**
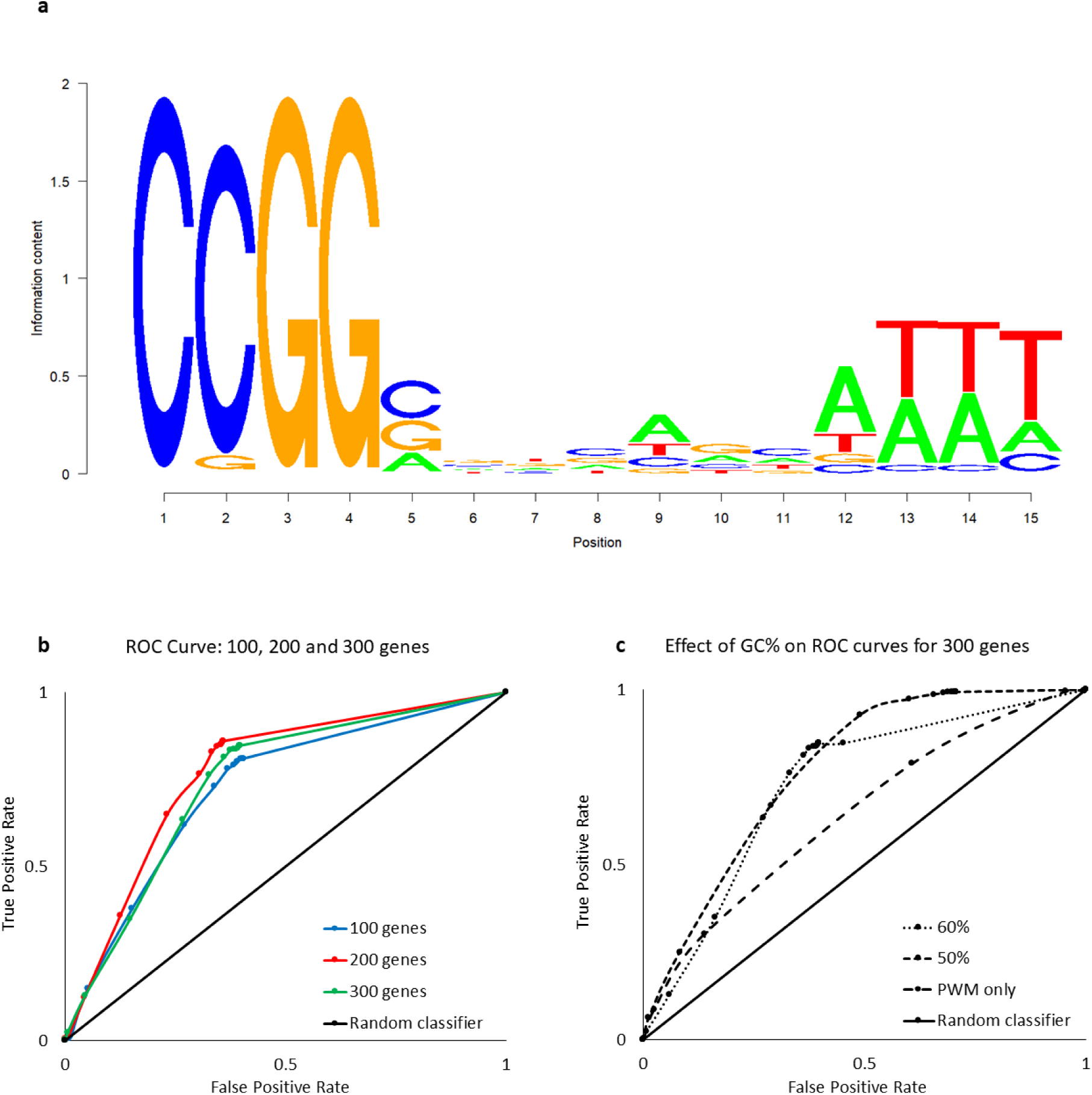
Establishing an MeCP2 position weight matrix threshold and GC%: Matrix-GC Procedure. **a** The MeCP2 sequence logo represents the conservation of sequence nucleotides for MeCP2. **b** ROC curves for 100, 200 and 300 genes establishing a preferential PWM score threshold. The AUC is 0.725, 0.7685, 0.7419, for 100, 200 and 300 genes respectively. **c** ROC curves for 300 genes evaluating the effects of DNA sequence GC content percentage. The AUC values are 0.7301, 0.7692 and 0.6351 for GC content percentages of 50%, 60%, and PWM only, respectively. The random classifier has an AUC of 0.50.

To verify that our PWM + GC% filter is effective in identifying genes bound by MeCP2, we apply the model to the top scored genes in the MeCP2 ChIP-Seq (Maunakea et al., 2013). We generate datasets of 100, 200 and 300 genes to determine if there was a preferential threshold at different ranking levels. The ROC curves represent the relationship between the true positive rate (TPR: sensitivity) and the false positive rate (FPR: 1-specificity) over a number of threshold values that can inform on the discriminating power of a performance model (Vihinen, 2012). Between 100, 200 and 300 genes, the three ROC curves are close to each other with similar AUC values of 0.725, 0.7685 and 0.7419 respectively (Fig. 1b, 1c). Considering the best trade-off between the TPR and FPR, we identify the ideal score threshold to be 65% for MeCP2 binding across the gene sets. Since MeCP2 has a higher binding potential for regions containing GC dinucleotide occurrence of ≥ 60% (Rube et al., 2016), we tested whether the PWM threshold score of 0.65 changes with different percentages of GC content. We combine the PWM filter with an additional GC content filter, varying the GC percentage threshold from 60%, to 50% and without filtering for GC content (PWM only). At 50% and 60% GC content (AUC = 0.7301 and 0.7692 respectively), the PWM + GC% filter performs better than both the random classifier and PWM filter alone. Despite 50% GC content having the largest AUC at the 65% PWM score threshold, 60% GC content offers a reduction in false positive rate by nearly half (50%: 0.657 vs. 60%: 0362 false positive rate). We confirm that a GC content of 60% is appropriate and in line with Rube and colleagues’ report (Rube et al., 2016) and use a PWM score of 65% and GC content of 60% for further analyses (Matrix-GC).

To further test the validity of our procedure, we consider the binding of *CDKL4* as a negative control. Cdkl4 has been reported to be downregulated in both MeCP2 overexpressing and null mice models with no detected MeCP2 binding in the promoter region (Chahrour et al., 2008). The current Matrix-GC procedure captures the same finding in mouse *Cdkl4* and human *CDKL4* orthologue promoter sequence. The use of these controls confirmed the validity of our method.

Then, we examine the MeCP2 binding potential according to Matrix-GC on gene sets associated with neuropsychiatric and neurological disorders (Fig. 2a, 2b). All neuropsychiatric datasets have at least 50% of genes putatively bound by MeCP2 through the Matrix-GC procedure. Neurological datasets show an overall lower average percentage of MeCP2-bound genes (55.95 %) compared to neuropsychiatric disorders (67.58%). These results suggest a major involvement of MeCP2 in neuropsychiatric pathologies, but it could be attributed to the lower numbers of genes present the neurological data sets. We also consider the genome and applied Matrix-GC to all the genes in the GRCh37/hg19 human reference genome (Fig. 2c). We report an average of 39.56% genes bound by MeCP2 *in silico* across the genome. The consistent higher percentage of bound genes in the disorder datasets compared to across the chromosomes is expected since they are brain disorders-related genes.

**Fig. 2.**
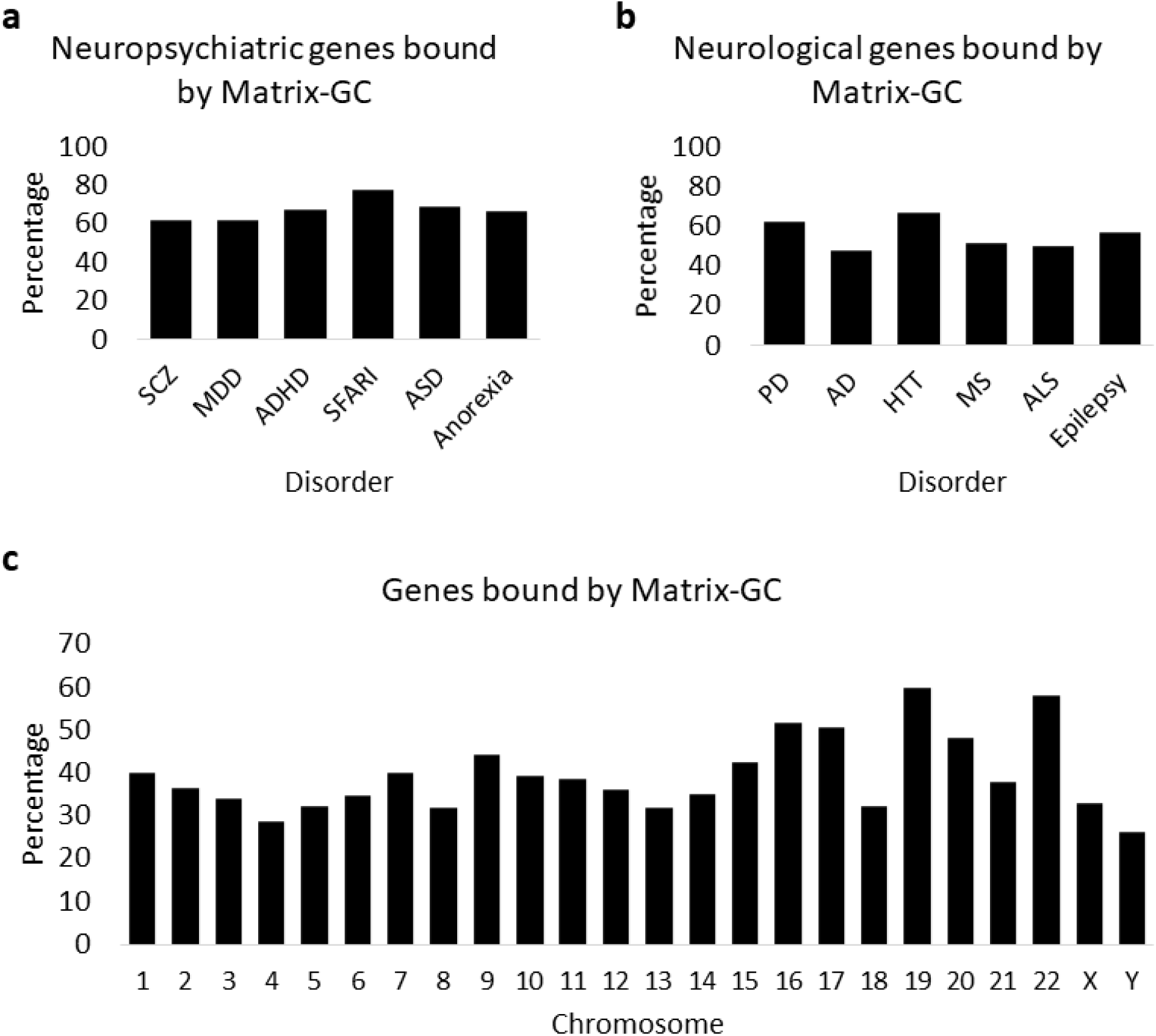
Genes bound by MeCP2 *in silico* in the genome and disorder datasets. **a** Percentage of genes bound by our Matrix-GC procedure in the neuropsychiatric datasets: schizophrenia (SCZ), major depressive disorder (MDD), attention deficit hyperactivity disorder (ADHD), Autism database (SFARI), Autism (ASD), and anorexia. **b** Percentage of genes bound by our Matrix-GC procedure in the neurological datasets: Parkinson’s disease (PD), Alzheimer’s disease (AD), Huntington’s disease (HTT), multiple sclerosis (MS), amyotrophic lateral sclerosis (ALS), and epilepsy. **c** Percentage of genes bound by our Matrix-GC procedure in the genome.

### Protein-Protein Interaction Network Analysis through Cytoscape

In order to uncover the molecular interaction guided by the candidate genes, we perform network analysis through Cytoscape using the StringAPP on genes filtered with the Matrix-GC procedure and we identify central proteins or nodes that are highly connected in each disorder.

From our PPI network, we identify hub proteins from neuropsychiatric datasets (65 proteins from SFARI, 34 from SCZ, 17 from ADHD, and 1 from MDD) with a node degree of ≥ 10 (Table 1). EP300, GRIN2B, NTRK3 and CNTNAP2 result as hub proteins from more than 1 dataset. Notably, EP300 is a hub protein in SFARI, SCZ and MDD with a degree of 45, 34 and 15, respectively. EP300 is a histone acetylase regulated indirectly by MeCP2 likely via MEF2C (Zweier et al., 2010). MECP2 itself is detected by our Matrix-GC procedure and is a hub protein according to PPI analysis (Supplementary Information File).

**Table 1.**
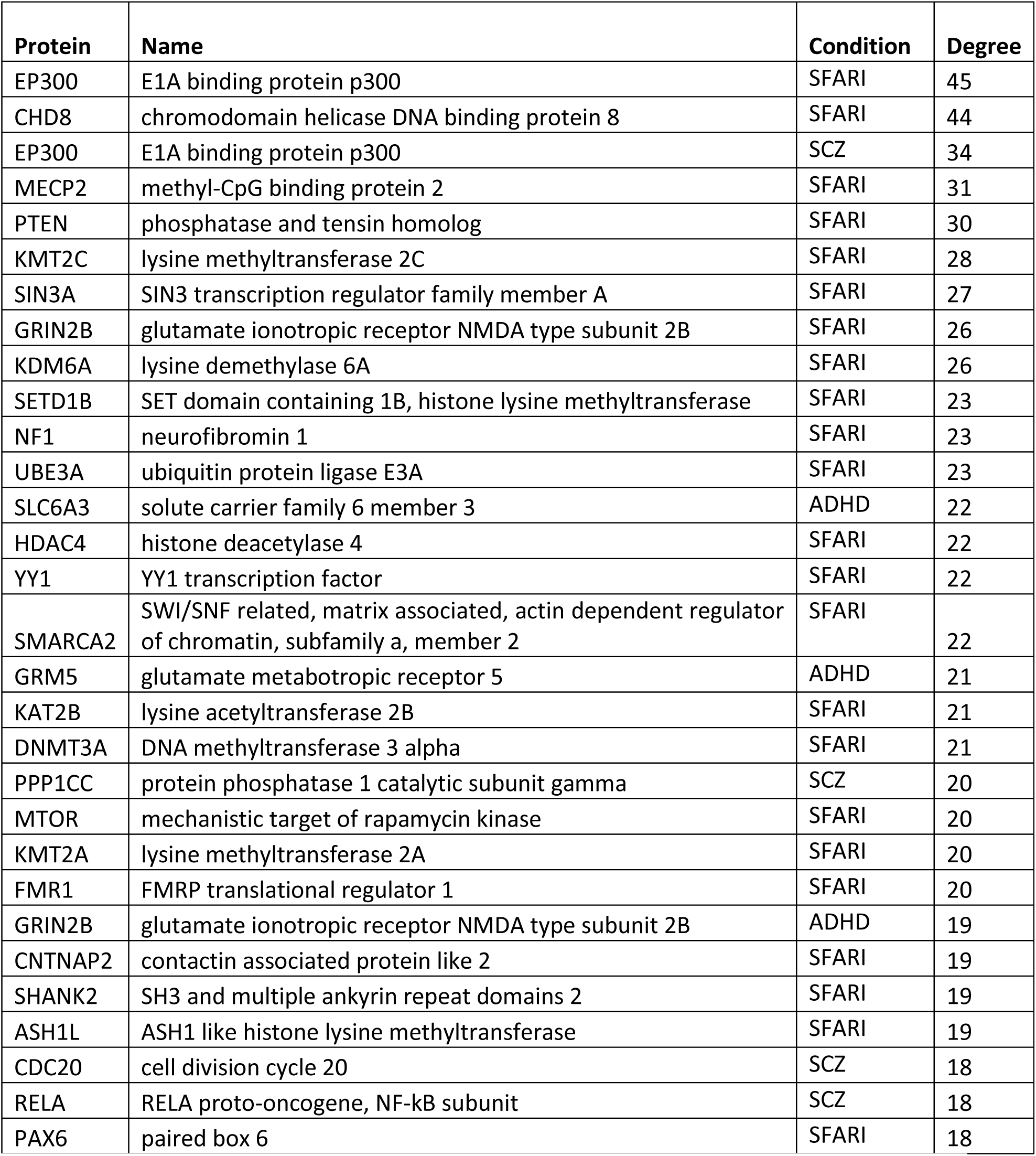
Top 30 hub proteins from protein-protein interaction analysis ranked by node degree.

The results of PPI analysis highlight proteins important in transcription regulation and neurotransmission.

### Enrichment Analysis Reveals Unexpected MeCP2 Influence in Neuropsychiatric and Neurological Disorders

In order to establish whether the candidate targets of MeCP2 are part of biological processes and pathways enriched in the disorder datasets, we carry out GO and pathway enrichment analysis in the data sets before and after the application of Matrix-GC.

We find ADHD, AD and SFARI to have significantly enriched GO Biological Process terms. SCZ processes were significant before and after the Matrix-GC procedure. SCZ terms are significant after Matrix-GC only. Epilepsy, MDD, MS, and PD show significant terms prior to Matrix-GC only.

Over-representation analysis shows that terms relating to neuronal growth, differentiation and nervous system development are significantly enriched in both ADHD and SCZ datasets. Additionally, ADHD genes show a significant enrichment in behaviour and learning, cell-cell communication, and catecholamine neurotransmission and metabolism. AD genes detected by the Matrix-GC procedure show significantly enriched terms related to amyloid protein regulation, metabolism, protein filaments and endocytosis. The SFARI dataset has the highest number of enriched terms before and after the Matrix-GC procedure, and the most significant terms relate to nucleic acid processes (Fig. 3).

**Fig. 3.**
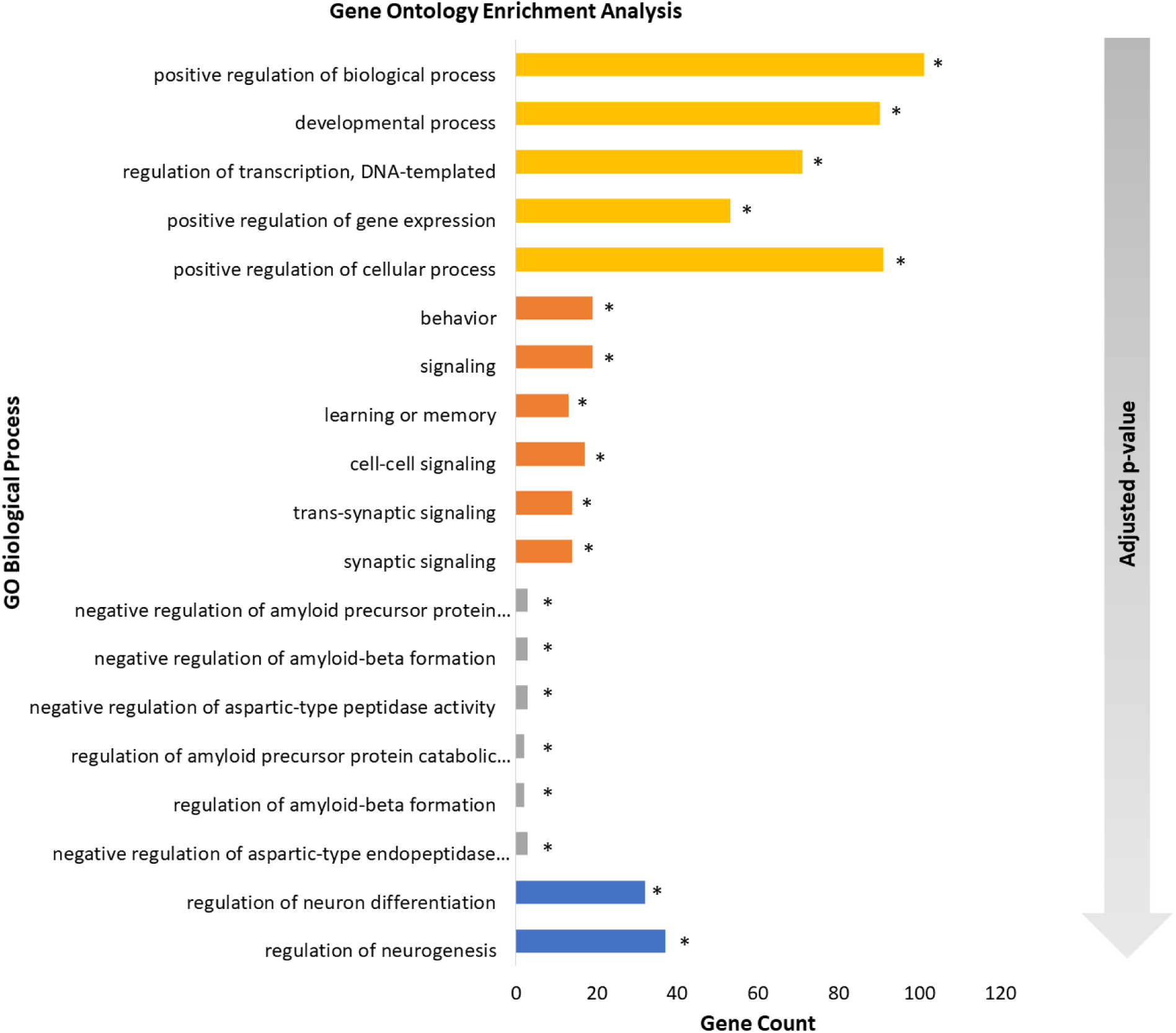
Gene Ontology enrichment analysis of neuropsychiatric and neurological disorders. Bar plot of the 5 most significant results from Gene Ontology enrichment analysis of each neuropsychiatric and neurological datasets after the Matrix-GC. The following disorders are represented: attention deficit hyperactivity disorder (orange), schizophrenia (blue), Alzheimer’s disease (grey), autism SFARI (yellow). The gradient arrow indicates that the adjusted p-values increase along the y-axis. * indicates terms that are also significant before Matrix-GC.

We then perform pathway analysis using Reactome to reveal significantly enriched pathways for neuropsychiatric and neurological disorders (Fig. 4). Only MDD, anorexia and epilepsy gene sets display enriched pathways solely before our Matrix-GC procedure. SCZ and ASD have no enriched pathways either before or after the Matrix-GC procedure. Conversely, PD pathways are significant after Matrix-GC only, and SFARI, AD, ADHD, ALS, HTT and MS have enriched pathways both before and after the procedure (Supplementary Information File).

**Fig. 4.**
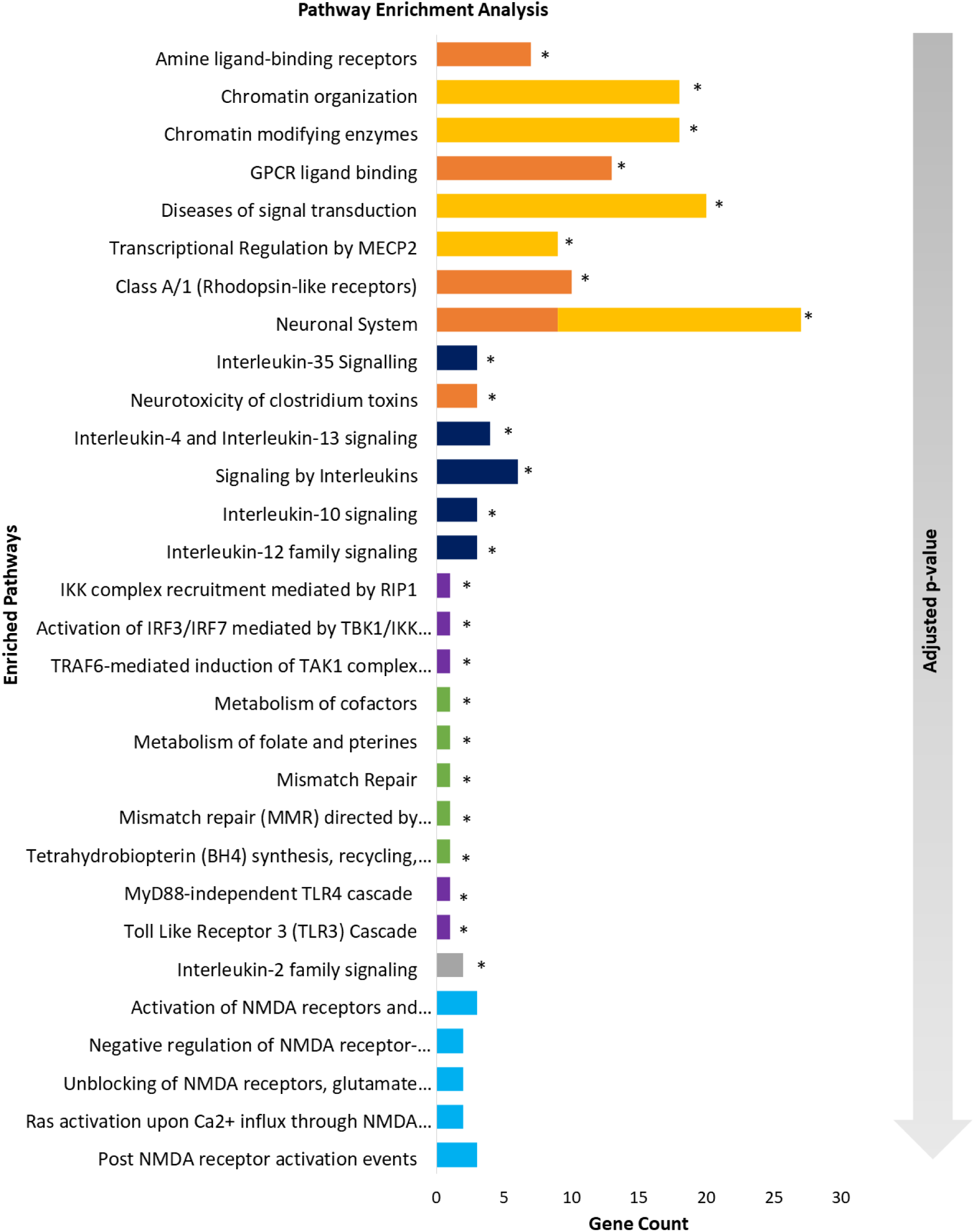
Pathway enrichment analysis of neuropsychiatric and neurological disorders. Bar plot of the 5 most significant results from pathway enrichment analysis of each neuropsychiatric and neurological datasets after Matrix-GC. The following disorders are represented: attention deficit hyperactivity disorder (orange), Alzheimer’s disease (grey), multiple sclerosis (dark blue), Parkinson’s disease (light blue) Huntington’s disease (green) and amyotrophic lateral sclerosis (purple), autism SFARI (yellow). The gradient arrow indicates that the adjusted p-values increase along the y-axis. * indicates pathways that are also significant before Matrix-GC.

**Fig. 4.**
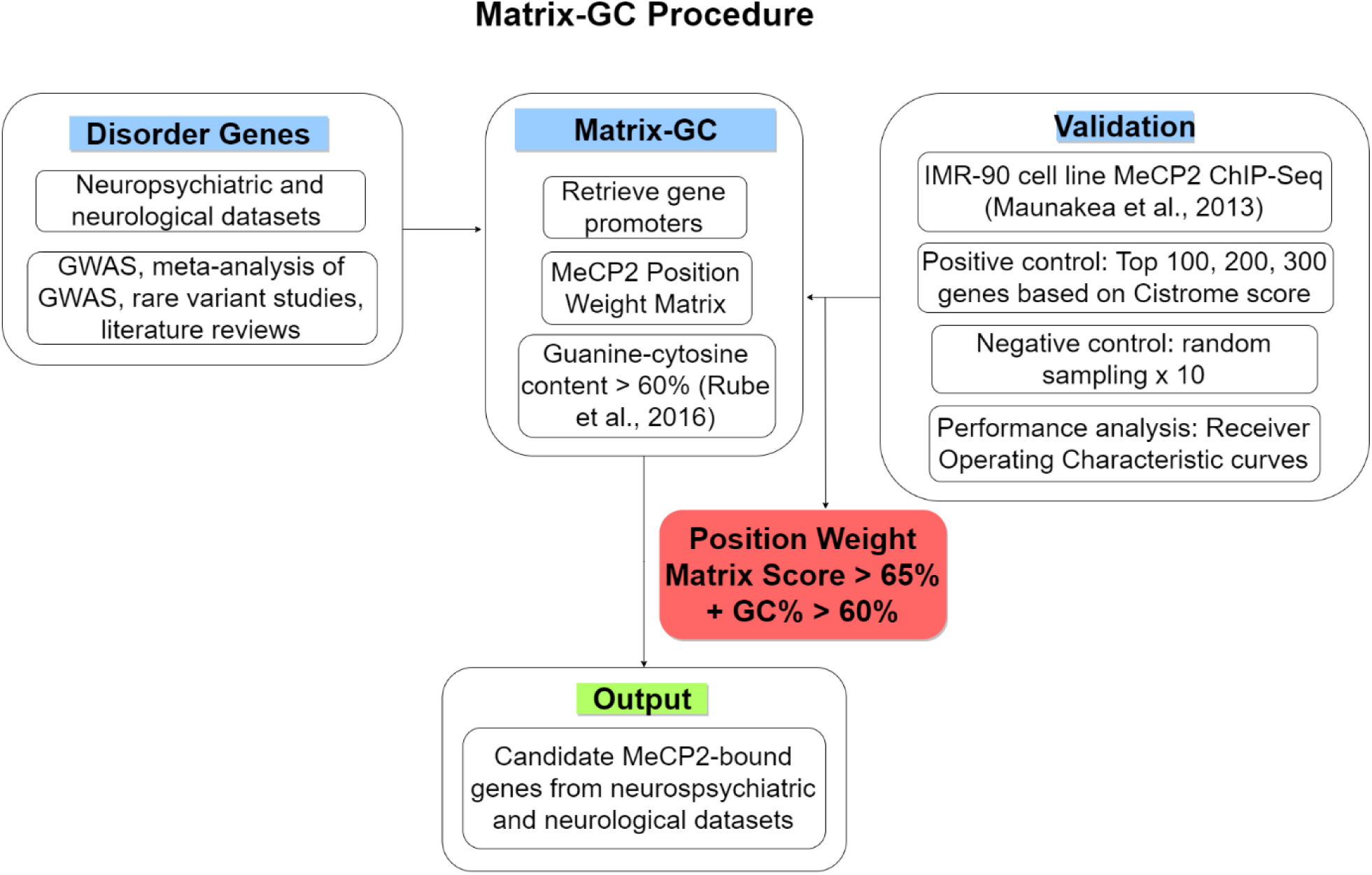
Overview of Matrix-GC procedure to detect MeCP2 transcription factor binding sites *in silico.* The Matrix-GC procedure is a combination of an MeCP2 position weight matrix and DNA sequence GC%, and was validated through positive and negative random sampling control using data from (Maunakea et al., 2013). The performance of Matrix-GC was evaluated through ROC curves. Matrix-GC was applied to the promoters of candidate genes across neurological and neuropsychiatric disorders to give an output list of putative genes bound by MeCP2 from each disorder.

Overall the most significantly enriched pathway is amine ligand-binding receptors in ADHD (adjusted p-value = 8.12 x 10^−8^). Glutamate and CREB related pathways are also enriched in ADHD, while SFARI has the highest number of overrepresented pathway associated with chromatin organisation, growth and neurotransmitter processes. AD and MS are significantly enriched for interleukin signalling pathways. ALS genes are enriched for Toll-like receptor (TLR) processes both before and after Matrix-GC. PD displays significant NMDA-related pathways. The use of control datasets (Supplementary Information File) confirms that these results are not by chance as most controls-related datasets do not appear in their corresponding disorder dataset. Only two pathways in the ALS control for Reactome analysis are significant and therefore we do not consider them among the positive results. Overall, we find the ADHD, AD and SFARI derived genes to be highly enriched for GO terms and Reactome pathways suggesting that several mechanisms controlled by MeCP2 are relevant in these disorders.

Overall, our analysis shows that candidate genes bound by MeCP2 in various disorders are enriched for terms and pathways broadly involving growth and development, neurotransmission and immunity.

## Discussion

In this study we consider the hypothesis that MeCP2, a protein mostly linked to Rett Syndrome, controls biological mechanisms and pathways in other brain disorders. We use a combination of bioinformatics tools, enrichment analysis and diseases-linked datasets, to identify potential MeCP2-modulated mechanisms. We show that MeCP2 binds to brain disorder candidate genes through Matrix-GC at a higher rate than across the genome (Figure 2). Our downstream enrichment analysis indicates that MeCP2 influences mechanisms mostly in Autism, ADHD, schizophrenia and Alzheimer’s disease. To our knowledge, this is the first study to outline a procedure for MeCP2 binding to identify binding sites *in silico* from disease-related gene sets, and it reveals common mechanisms between different brain disorders.

The role of MeCP2 as a regulator of brain development and function is confirmed by multiple publications in both patients and animal models reporting *MECP2* mutations in brain disorders. Most MeCP2 disease-causing mutations result in a partial or total loss of function, although there are reports of MeCP2 duplication with individuals presenting with autistic and neurospychiatric traits (Ramocki et al., 2009). A recent study in transgenic primates overexpressing MeCP2 (Liu et al., 2016) showed that compared to wild conspecifics, transgenic macaques had reduced interaction times in groups and pairs and showed larger variability in cognitive function tests. Overexpression of MeCP2 is related to ASD (Peters et al., 2013), reflecting in stereotyped behaviours in patients and rodent models.

Despite the strong association between ASD and MeCP2, enrichment and network analysis based only on analysis of common variants do not identify terms or pathways in ASD GWAS dataset (Grove et al., 2019). These results are surprising, although we find several enriched mechanisms in ASD when considering the larger SFARI dataset which takes into account all genes associated with autism. It is possible that MeCP2 might play a role in coordinating functional connectivity through controlling delicate neurotransmitter balance in ASD as we see in other neuropsychiatric disorders (Ausió et al., 2014).

In SCZ, mutations in the MeCP2 AT-domains have been identified in one patient (McCarthy et al., 2014), and one of the endophenotypes: reduced connectivity, has been reported in both SCZ and RTT. MeCP2 binding to GABAergic genes is increased in mice with prenatal restraint stress, resulting in psychotic-like phenotype similar to SCZ. MEF2C is a transcription factor that binds to the MeCP2 promoter and controls MeCP2 expression (Zweier et al., 2010). MEF2C mutations affect MeCP2 function and this is observed in epilepsy and ADHD (Paciorkowski et al., 2013). It is also noteworthy to stress that MeCP2 mutations can be present without RTT diagnosis, and this is seen in individuals with neurodevelopmental deficits, OCD and ADHD (Suter et al., 2014).

The influence of MeCP2 in epilepsy is confirmed by seizures reported in RTT and by altered MeCP2 expression (Tao et al., 2012) reported in epilepsy. In PD, under-expression of MeCP2 in dopaminergic neurons of the substantia nigra, contributes to motor deficits; the same neurons alter electrochemical and neuron development in RTT mice throughout development and into adulthood (Gantz et al., 2011). Blood transcriptomics has also shown that MeCP2 is a differentially expressed gene in PD (Calligaris et al., 2015). Lead-exposure mediated increase of AD-related protein is associated with a decrease of MeCP2 and DNA methyltransferases in cerebral cortex (Eid et al., 2016). In ALS, mutations in *FUS* disrupts MeCP2 expression through alterations in mRNA splicing and stability leading to lower levels of MeCP2 (Coady and Manley, 2015). Similar to MS, RTT displays features that are hallmarks of autoimmune disorders (De Felice et al., 2016). Mir124 is downregulated in experimental autoimmune encephalomyelitis (EAE) models of MS (Soreq and Wolf, 2011). Mir124 downregulates MeCP2’s recruitment during development to promote neuron formation by supressing non-neuronal genes (Visvanathan et al., 2007). Recent work shows that non-psychoactive cannabinoids -a possible treatment for MS (Baker et al., 2000)-ameliorates behavioural alterations and partial motor deficits in *Mecp2* mutant mice (Vigli et al., 2018).

The broad scope of MeCP2 in regulating neuronal activity and nervous system function justifies the wide clinical presentation of RTT and the overlapping of symptoms with other brain disorders (Ausió et al., 2014). It is plausible that MeCP2 controls specific cellular mechanisms in several disorders considering its role in GABAergic, serotonergic, dopaminergic transmission, as well as in the function of glutamatergic synapses. In addition, MeCP2 controls the expression of Brain-derived Neurotrofic Factor (BDNF), a neurotrophin involved in brain development, function and connectivity (Chen et al., 2003). BDNF-related mechanisms are dysregulated as a result of MeCP2 mutations or altered expression levels. It is possible that MeCP2 controls brain disorders through BDNF. BDNF altered expression has been detected in several disorders, including NDD, MDD, and anxiety. DNA methylation within the BDNF promoter is altered in bipolar disorder. BDNF is reported to be increased in SCZ post mortem brains. In eating disorders, BDNF is reported to play a role in feeding behaviour (Autry and Monteggia, 2012).

Overall our analysis reveals three main mechanisms controlled by MeCP2 in different disorders: neuronal transmission, immune-related pathways and processes for growth and development.

As part of neuronal transmission, we find enrichment in dopaminergic and glutamatergic related terms and pathways in ADHD, SFARI, and PD, which is reinforced by hub proteins related to DA and glutamate receptors.

Dopaminergic dysregulation in RTT patients have been observed through reductions in DA or its metabolite, homovanillic acid in plasma, cerebral spinal fluid, and postmortem brain tissue (Riederer et al., 1985; Lekman et al., 1989; Zoghbi et al., 1989). This dysregulation leads to dyskinesia, hand stereotypies and rigidity: all symptoms found in RTT. Recently, Wong and colleagues showed that *Mecp2* -/y mice exhibit similar brain pathology detected in human observations where D2 receptor and DA transporter densities were compared (Wong et al., 2018). Alteration in dopamine transmission is a feature of several neurological disorders, notably PD, but also AD, MS and HTT (Wolfe et al., 1990; Dobryakova et al., 2015). It is known that DA tracts also play a role in the aetiology of several neuropsychiatric diseases such as ASD, SCZ and ADHD (Volkow et al., 2009; Money and Stanwood, 2013) where many DA genes are disrupted. There is a strong association between ADHD and the DA mesolimbic pathway, and there are deficits in reward and motivation leading to inattention and impulsivity (Volkow et al., 2009). We show that NMDA receptor-related pathways are enriched in the ADHD and PD dataset which suggests MeCP2’s influence in altering glutamatergic signalling. Additionally, several hub genes present in SCZ relate to DA and glutamate receptors which is expected since abnormal signalling of both neurotransmitters is observed in this disorder (Howes et al., 2015).

Increased levels of glutamate, localised in cerebrospinal fluid (Lappalainen and Riikonen, 1996) are observed in RTT. *Mecp2*-null microglia are shown to cause damage to dendrites and synapses through increased release of glutamate (Maezawa and Jin, 2010). Additionally, glutamatergic synapses are regulated by MeCP2. A recent study showed that MeCP2 deficiency in glutamatergic neurons results in neurobehavioral deficits in mice and is ameliorated in female *Mecp2*-heterozygous mice when MeCP2 is restored (Meng et al., 2016). These results suggest that MeCP2 influence in dopaminergic and glutamatergic systems has functional and behavioural consequences in several brain disorders.

Immune related pathways and processes, namely interleukin signalling through various members of the interleukin (IL) family, are relevant because of the interplay between neuronal and immune function in several disorders. AD, ALS and MS are enriched for interleukin and TLR signalling pathways. Several cytokines are involved in the development of the central nervous system and are responsible for positively regulating key processes in development and maintenance of neuronal circuits (Yirmiya and Goshen, 2011). Furthermore, chronic inflammatory and depression modulate neurotransmission and behaviour (Felger and Lotrich, 2013). Inflammation and immune response are also related to dopaminergic transmission. DA plays a neuroregulatory role in controlling immune response in the the body. DA has been observed in murine and human immune cells; human lymphocytes express all DA receptor subtypes. During an immune response, DA regulates leukocyte cytokine production without any discrimination towards a proinflammatory or anti-inflammatory phenotype (Arreola et al., 2016). In clonal T-lymphocyte cultures, there are differentially expressed genes that implicate MeCP2’s involvement in regulating immune cells (Delgado et al., 2006). It would be no surprise that MeCP2 might play a role in coordinating these events as there are many immunological and neurotransmission changes in RTT. Cytokines are key players in the CNS, acting as trophic factors but can also augment neural homeostasis and lead to neural insults in neurological diseases such as PD, AD, MS. Altered immunity has also been reported in neuropsychiatric disorders such as SCZ, depression and ASD (Kerr et al., 2005; Khandaker et al., 2015). Interestingly, we observe a single GO term (GO:0002292, FDR= 4.13 x 10^−2^) and no pathways related to the immune system in the SFARI dataset.

Growth and developmental processes appear across the different datasets in the enrichment analysis with terms and pathways related to cell cycle and proliferation.This is expected, given that MeCP2 is an epigenetic modifier in cancer (Lengauer and Issa, 1998). GO enrichment analysis shows terms in SCZ and SFARI involved in cell and neuronal development, while network analysis shows hub proteins involved in chromatin remodelling, cell cycle and RNA processing. Taken together, these results suggest that MeCP2 exerts influence in early development in SCF and ASD. Similarly, ADHD is enriched for pathways related to neurotrophic factor signalling which mediates neuronal proliferation and maturation (Ghosh et al., 1994).

Overall, this is the first study to show a contribution of MeCP2 to mechanisms linked to neuronal transmission, neuro-immunity and development across several and unexpected brain disorders. These findings contribute understanding to the neurobiology of RTT and other pathologies and may suggest common therapeutic targets across disorders.

## Methods

### Establishing MeCP2 Binding

#### Position Weight Matrix

A PWM represents the motifs most likely to be bound by a TF allowing for redundancy in binding site specificity. PWMs are generated by combining experimental data for TFs binding DNA sequences (Stormo et al., 1986) and provides an *in silico* approach to predict binding sites for a specific TF. One limit of PWMs is the discrimination of true binding sites from non-binding sites as each nucleotide probability at nucleotide position is calculated independently from its neighbour (Pan and Phan, 2008). PWM sequence motifs can occur repeatedly across the genome, representing true binding sites or sites bound by chance. This poses a challenge in TF binding site analysis and the traditional PWM-only filter require additional information to improve discrimination.

We used a PFM for MeCP2 from the Cistrome database (http://cistrome.org) based on SELEX experiments carried out by Klose and colleagues (Klose et al., 2005). We used the Biostrings package in RStudio version 1.1.463, to convert the MeCP2 PFM into a PWM that could be used to identify the MeCP2 binding motif along a sequence of DNA (Fig. 1a). Klose and colleagues identified preferred sequences for MeCP2 binding through methyl-SELEX (Klose et al., 2005) and validated with genes known to be bound by MeCP2, *Bdnf* and *Dlx6* in the promoter and core gene regions.

### PWM Scanning

#### Positive and Negative Chip-Seq Control Datasets

We retrieved MeCP2 potential target genes from ChIP-Seq data Cistrome Data Browser (http://cistrome.org/db/). We used ChIP-Seq data from a study by Maunakea and colleagues (Cistrome ID 34392) (Maunakea et al., 2013). We used IMR-90 foetal lung myofibroblast data from this study as our positive control dataset. Putative target genes on Cistrome are already scored by the Binding and Expression Target Analysis package indicating the regulatory potential as a putative target (Wang et al., 2013).

For our positive controls, we generated sequence datasets for the top 100, 200 and 300 genes bound by MeCP2 from the IMR-90 foetal lung myofibroblast ChIP-Seq data, ranked by Cistrome BETA scoring. For our negative controls, we randomly selected and size-matched genes from the same ChIP-Seq data with a score of 0. We define the promoter sequences as being 1000bp upstream of the transcription start site and we retrieved these promoters in RStudio from the UCSC Genome Browser (https://genome.ucsc.edu/) using the GRCh37/hg19 human reference genome. We tested each sequence for the presence of the MeCP2 PWM. For every PWM match, a score is given from 1-100%. This score represents how the sequence being matched is different from a random sequence.

Guanine-Cytosine nucleotide content (GC%) was previously established to be important in MeCP2 binding *in vivo* (Rube et al., 2016). For every PWM match, we generated a sequence to include the 15bp PWM match sequence and 100bp flanking sequences, and we calculate the GC for these 215bp sequences.

#### Receiver Operating Characteristics Curve

In order to determine the ideal PWM threshold for MeCP2 motif binding, we graphed a ROC for all datasets. We set the minimum PWM score at 5% and stratified results based on PWM scores at increasing increments of 5%. For bootstrapping analysis, we generated 10 random samples of 100, 200 and 300 negative control genes. Taking the average values, we plotted the ROC curve alongside the positive controls. Additionally, we evaluated ROC curves at various sequence GC%. The AUC was calculated as a measure of performance relative to a random classifier (AUC = 0.5) that is represented by the line x = y (Figure 2).

#### Brain Disorders Dataset Collection

For the analysis of MeCP2 interaction in different disorders, we used datasets for different neuropsychiatric and neurological disorders gathered from multiple studies including GWAS, meta-analysis of GWAS, and literature reviews (Table 3). We exclude the extended major histocompatibility complex region from the SCZ dataset as it spans several hundred genes and potentially introduces noise into our analysis (Pardiñas et al., 2018). Additionally, the previous comprehensive gene map was described in 2004 and excluded RNA genes (Horton et al., 2004). For the SFARI dataset (Abrahams et al., 2013) Gene Scoring - which assesses the strength of evidence presented for a candidate ASD gene - we considered categories S (syndromic), 1 (high confidence genes) and 2 (strong candidate genes).

**Table 3.** List of neuropsychiatric and neurological diseases used in the present study.

| Disorder | Source | Journal | Data Type |
| --- | --- | --- | --- |
| **Neuropsychiatric** |  |  |  |
| Autism (ASD) | Grove et al., 2019 | Nature Genetics | Meta-analysis of GWAS |
| Autism (SFARI) | SFARI GENE |  | Database |
| Schizophrenia GWAS (SCZ) | Pardiñas et al., 2018 | Nature Genetics | GWAS |
| Attention deficit hyperactivity disorder (ADHD) | Hayman and Fernandez, 2018 | Frontiers in Psychiatry | Review of GWAS, copy number variant, and candidate gene association studies |
| Major depressive disorder (MDD) | Howard et al., 2019 | Nature Neuroscience | Meta-analysis of GWAS |
| Anorexia | Duncan et al., 2017 | American Journal of Psychiatry | GWAS |
| **Neurological** |  |  |  |
| Parkinson's disease (PD) | Chang et al., 2017 | Nature Genetics | Meta-analysis of GWAS |
| Alzheimer's disease (AD) | Lambert et al., 2013 | Nature Genetics | Meta-analysis of GWAS |
| Huntington's disease (HTT) | Moss et al., 2017 | The Lancet Neurology | GWAS |
| Multiple sclerosis (MS) | Bashinskaya et al., 2015 | Human Genetics | GWAS review |
| Amyotrophic lateral sclerosis (ALS) | van Rheenen et al., 2016 | Nature Genetics | GWAS |
| Epilepsy | Abou-Khalil et al., 2018 | Nature Communication | Meta-analysis of GWAS |

For SCZ associated genes, we used genes identified by GWAS studies. Rare variants are also implicated in SCZ aetiology, however only few reliable candidates have been identified until now: studies looking at rare single nucleotide variants require larger samples to meet power requirements as rare risk loci often occur at low frequencies. The study of CNVs can potentially identify single genes which are rare variants for SCZ, but CNVs at implicated loci can span several genes and brings into question which gene contributes to the phenotype. At the moment the few candidate rare variants in SCZ have been confirmed with sequencing. Neurexin 1 is a well-known CNV in SCZ (Marshall et al., 2017) and is also largely associated with ASD (Kim et al., 2008). To date, SETD1A is the only genome-wide significant rare variant discovered by whole exome sequencing (Singh et al., 2016). It proves difficult to find SCZ-exclusive rare variants and to this effect, we do not consider SCZ CNVs in our study.

### Enrichment and Network Analyses

We employed Gene Ontology and pathway overrepresentation analysis to define functional aspects of the disease gene datasets and identify any terms or pathways that were significantly enriched in these datasets. We carried out Gene Ontology enrichment analysis using GOrilla (http://cbl-gorilla.cs.technion.ac.il/). We discovered over-represented Gene ontology terms by analysing disorder genes versus the background set which ignores ranking. We consider the background set of genes as being the entire genome minus the genes of interest. To perform pathway enrichment analysis within the various gene sets, we input genes as Ensembl identifiers with a selected p-value threshold of 0.05. For pathways analysis we used the ReactomePA package from Bioconductor (https://bioconductor.org/packages/release/bioc/html/ReactomePA.html) and we input Entrez identifiers into the function call enrichPathway and selected a p-value cut-off value of 0.05, controlling for false discovery rate (“fdr”). For network analysis we used Cytoscape application version 3.7.1 and we visualised proteomics data from STRING (http://string-db.org) using the stringAPP plugin version 1.4.2 (http://apps.cytoscape.org/apps/stringapp). We input Ensembl identifiers to identify any protein-protein interactions, either directly or indirectly. We applied a confidence cut-off value of 0.4 with 0 additional interactors and applied the NetworkAnalyzer function in the StringAPP for network analysis and the edges were treated as undirected. We designated hub proteins as having a node degree of ≥ 10.

One possible caveat in using this approach is the different number of genes present in the datasets considered, and the limited number of genes in several disorders (anorexia, epilepsy, HTT) as these datasets have lower statistical power when it comes to enrichment and network analysis. Hence the network and enrichment component of the study has a natural bias toward the disorders with more genes. To minimise the intrinsic biases of enrichment and network analysis we used random controlled datasets with the same number of genes used as input. For control enrichment and network analyses, we generated 10 random samples of genes from the genome and run them through the enriched analysis. The size of the controls datasets was the same as the corresponding input for each disorder.

## Acknowledgements

The work had a partial contribution by EIT Health (19069) grant to D.T. S.B.’s fees are funded by SFI (16/IA/4443) grant to P.G.

## Author contributions

D.T., N.M.R and S.B. designed the research and S.B. conducted the bioinformatics methods. S.B., N.M.R, and D.T prepared the manuscript. P.G. advised on statistical methods and assisted in preparing the manuscript. A.C. participated in the selection of the datasets and discussing the results.

## Additional information

## Abbreviations

AD: Alzheimer’s Disease
ADHD: Attention Deficit Hyperactivity Disorder
ALS: Amyotrophic Lateral Sclerosis
ASD: Autism Spectrum Disorder
AUC: Area Under the Curve
BDNF: Brain-Derived Neurotrophic Factor
CDLK4: Cyclin-Dependant Kinase-Like 4
CNV: Copy Number Variant
DA: Dopamine
GWAS: Genome-Wide Association Study
HTT: Huntington’s Disease
MDD: Major Depressive Disorder
MeCP2: Methyl-CpG Binding Protein 2
MEF2C: Myocyte Enhancer Factor 2C
miRNA: Micro RNA
MS: Multiple Sclerosis
NDD: Neurodevelopmental
OCD: Obsessive Compulsive Disorder
PD: Parkinson’s Disease
PPI: Protein-Protein Interaction
PFM: Position Frequency Matrix
PWM: Position Weight Matrix
RTT: Rett Syndrome
ROC: Receiver Operated Characteristics
SCZ: Schizophrenia
TF: Transcription Factor

## Data availability

Additional data supporting the results of this study are available within this article and in the Supplementary Information file, and from the corresponding author upon reasonable request.

## Code availability

The detailed procedure and R scripts used in the analysis supporting the findings of this study are available from the corresponding author upon reasonable request.

